# Assessing the role of trypsin in quantitative plasma- and single-cell proteomics towards clinical application

**DOI:** 10.1101/2023.05.26.542422

**Authors:** Jakob Woessmann, Valdemaras Petrosius, Nil Üresin, David Kotol, Pedro Aragon-Fernandez, Andreas Hober, Ulrich auf dem Keller, Fredrik Edfors, Erwin M. Schoof

**Author notes:** Contributed equally to this work (Fredrik Edfors, Erwin M. Schoof).

## Abstract

Mass spectrometry-based bottom-up proteomics is rapidly evolving and routinely applied in large biomedical studies. Proteases are a central component of every bottom-up proteomics experiment, digesting proteins into peptides. Trypsin has been the most widely applied protease in proteomics, due to its characteristics. With ever-larger cohort sizes and possible future clinical application of mass spectrometry-based proteomics, the technical impact of trypsin becomes increasingly relevant. To assess possible biases introduced by trypsin digestion, we evaluated the impact of eight commercially available trypsins in a variety of bottom-up proteomics experiments, and across a range of protease concentrations and storage times. To investigate the universal impact of these technical attributes, we included bulk HeLa-cell lysate, human plasma and single HEK293 cells, which were analyzed over a range of Selected Reaction Monitoring (SRM), Data-Independent Acquisition (DIA), and Data-Dependent Acquisition (DDA) instrument methods on three LC-MS instruments. Quantification methods employed encompassed both label-free approaches and absolute quantification utilizing spike-in heavy-labeled recombinant protein fragment standards. Based on this extensive dataset, we report variations between commercial trypsins, their source, as well as their concentration. Furthermore, we provide suggestions on the handling of trypsin in large scale studies.

## Introduction

The field of mass spectrometry (MS) driven proteomics has had major impact on biomedical research. Its application in precision medicine efforts has become more prominent in the past years, ranging from plasma proteome analysis of liquid biopsies to single-cell spatial proteomics [1][2]. During this process, increasingly larger cohorts are being analyzed, new techniques developed and standardized, while in unison moving the field closer towards the realms of clinical application [3]–[9]. Here, especially high reproducibility and robustness are key to drive MS forward into clinical practice and impactful biological discoveries.

Due to their systemic representation and ease of access, biofluid-based biopsies such as serum and plasma are attractive target samples for predictive biomarker panels that can be translated into clinical tests. However, for global MS-driven analysis, the complexity of the plasma proteome with its estimated 10^10^ dynamic range provides great challenges [10]. Here depletion methods and fractionation of high-abundant proteins can be applied to access low-abundant proteins [2], [11].

Another technology of increasing interest within the realm of MS is single-cell proteomics by MS (scp-MS). It has emerged as a promising novel technique, which is moving away from being predominantly a technical exercise to having real-life biomedical implications [12], [13]–[15]. Here the extremely limited input material and low abundance of proteins in an individual cell provides major technical challenges, as summarized in [12], [13], [16]–[18]. These challenges somewhat oppose the hurdles that the technology is facing in the plasma proteome field.

Both plasma as well as scp-MS have seen tremendous advances with the application of different MS acquisition techniques, labelling techniques, and sample preparation [8], [19]. When it comes to sample preparation, the field of plasma proteomics has seen major advancements in recent years, putting large efforts into optimizing and standardizing preparation techniques [7], [20]–[22]. After a plethora of individual efforts by leading labs, the field of sc proteomics published its first joint recommendations in early 2023 [23]. Among the main challenges, sample loss due to surface is a critical factor, which is of much greater importance in scp-MS than e.g. high load plasma samples. This obviates the need for avoiding commonly used lab approaches such as e.g. desalting, especially in the case of label-free quantification (LFQ) approaches. Therefore, denaturing conditions must be adjusted and different MS-compatible lysis reagents such as TFE, DDM, DMSO etc. have been used [24]– [26].

However, irrespective of sample type, the vast majority of global, data-driven LC-MS proteomics rely on a bottom-up approach, where proteases are the driving force to turn denatured proteins into peptides. Trypsin has been used predominantly for this task even though other proteases have been explored to increase proteome depth and quantification [27]–[31]. Trypsin is a serine protease that is produced in the pancreas of most vertebrates. It has a high specificity for cleavage C-terminal of Lysine and Arginine [32]. Trypsin generates peptides that are favorable for electrospray ionization (ESI) MS approaches, combined with online C18 reverse phase liquid chromatography [33]. Furthermore, tryptic peptides benefit from widely used CID and higher-energy HCD fragmentation methods [34], [28]. However, as an enzyme, trypsin is susceptible to digestion and reconstitution conditions which can impact enzymatic activity and digestion efficiency [20], [35]–[37]. Moreover, it has also been shown that the choice of trypsin can impact the identified number of peptides and the number of missed cleavages [38], [39]. There have been a number of publications reaching back a decade comparing trypsin performance of different vendors. In general, a variation between trypsins of different vendors has been reported [40], [41]. Others also looked into the differences between bovine and porcine trypsins [42]. In the field of scp-MS significant differences in the number of identified proteins depending on the protease concentration and two commercially available trypsins has been shown [25].

Today there is a large variety of trypsins by different vendors on the market that are mainly recombinantly expressed or extracted from porcine pancreatic extract. Some of them are further modified to decrease autolytic degradation or have been treated with tosyl phenylalanyl chloromethyl ketone (TPCK) to reduce chymotrypsin activity. With an ever-increasing number of biological samples included in clinical cohorts, the relevance of exploring to what extent trypsin might act as a technical variable, and as a consequence, introducing batch effects becomes more important. Here the effect of trypsin vendors or storage time on batch effects must be addressed. This becomes even more relevant once the field transitions into real-time clinical testing, differing vastly from a well-planned analysis of retrospective clinical cohorts. Finally, another relevant aspect is the impact of trypsin concentration, especially in terms of peptide quantification, and when comparing multiple datasets. In scp-MS trypsin has been shown to impact the number of peptides identified [25]. Here the impact of trypsin on reaching the lower abundant proteins is relatively unexplored.

To address the impact of trypsin with regard to future clinical applications of bottom-up proteomics, we here investigate to what extent we can detect experimental variations due to trypsin source, vendor and storage time (**Figure 1**). Eight trypsins from four vendors were assessed in HeLa bulk cell lysate, human plasma and single HEK293 cells. To assess the impact on different experimental setups, we evaluated data-dependent acquisition (DDA), data-independent acquisition (DIA) and selected reaction monitoring (SRM) to represent all commonly used data acquisition approaches. Furthermore, LFQ and absolute quantification were explored by incorporating heavy labeled standards for the absolute quantification of 122 plasma proteins with SRM. Combined, we aim to provide an overview of the influence of trypsin on reproducibility and quantification when transitioning further into the clinical application of MS driven proteomics.

**Figure 1.**
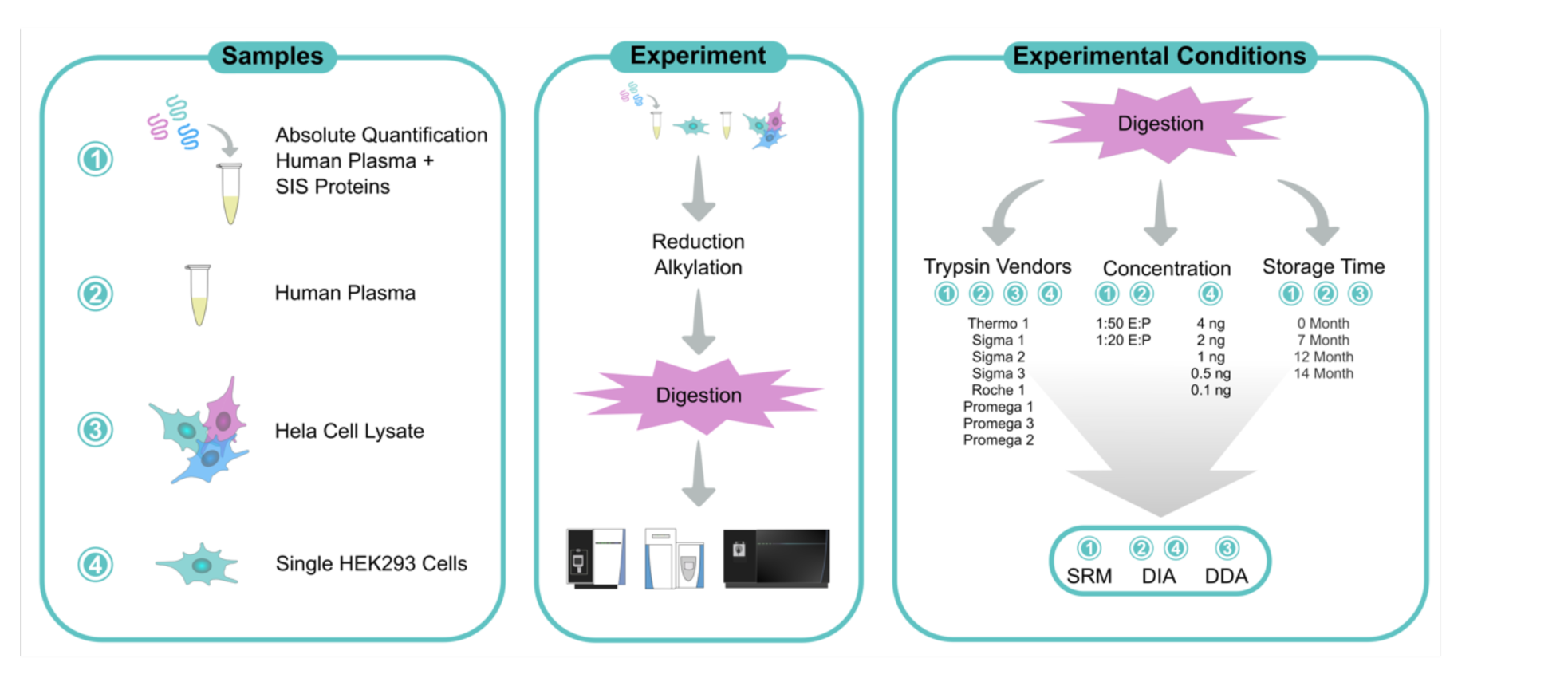
Overview of the experimental setup, showing all sample types, their respective LC-MS analysis methods, the specific trypsins used, trypsin concentrations and storage times that were investigated.

## Material and Methods

### Trypsin

The eight trypsins (**Table 1**) which were compared on HeLa cell lysate and human plasma were received within seven days and stored according to vendors recommendation immediately after arrival. Trypsins were reconstituted according to vendors instructions and samples were digested two weeks after trypsins were received. To assess storage time related impacts of trypsin, “Thermo 1“ was stored lyophilized at four timepoints at recommended temperature without interruption. “Thermo 1“ was stored for 0, 7, 12, and 14 months prior to sample preparation. Promega 1-3, “Sigma 3” and “Thermo 1” were compared for single cell experiments. They were stored at recommended temperature for up to 4 weeks prior to reconstitution and use.

**Table 1.** Summary of compared Trypsins.

| ID | Vendor | Product No. | Source |
| --- | --- | --- | --- |
| Promega 1 | Promega | V5111 | Porcine pancreatic extracts |
| Promega 2 | Promega | V5280 | Porcine pancreatic extracts |
| Promega 3 | Promega | VA9000 | Expressed in *pichia pastoris* |
| Roche 1 | Roche | 3708985001 | Expressed in *pichia pastoris* |
| Sigma 1 | Sigma-Aldrich | EMS0006 | Expressed in *pichia pastoris* |
| Sigma 2 | Sigma-Aldrich | EMS0007 | Expressed in *pichia pastoris* |
| Sigma 3 | Sigma-Aldrich | EMS0004 | Expressed in *pichia pastoris* |
| Thermo 1 | Thermo Fisher Scientific | 90058 | Porcine pancreatic extracts |

### HeLa cell lysate

HeLa-cells were cultured in Dulbecco’s Modified Eagle Medium, supplemented with 10% fetal bovine serum and 1% non-essential amino acids at 37 °C, 5 % CO_2_ in a humidified environment. Cells were washed in ice cold PBS, resuspended in 1x PBS, 1 % SDC and heat treated at 96 °C for 10 min. Cells suspension was aspirated and dispensed though 23G needle 10 times followed by sonication. Cell lysate was centrifuged at 17000 x g for 30 min and the protein concentration of the supernatant was measured using the Bio-Rad protein assay. Cell lysate was stored at -80 °C till further sample preparation.

### Human Plasma

Plasma (K_2_-EDTA) from five healthy individuals was pooled and stored at -80 °C (three male and two female). This plasma pool was used for labeled- and label free experiments. The samples were taken after informed consent by each individual as approved by the regional ethics board in Stockholm, and conducted in accordance with the principles set out in the Declaration of Helsinki.

### SIS-PrEST preparation

Stable isotope labeled protein epitope signature tags (SIS-PrESTs, ^13^C and ^15^N labeled) are recombinantly expressed protein standards that are up to 149 amino acids (AA) long [43]. They represent a unique part of endogenous human proteins and can be digested with different proteases [30]. SIS-PrESTs for 201 plasma protein targets were prepared within the Human Protein Atlas as previously described [44]. SIS-PrESTs were absolute quantified as previously described [45]. They were adjusted to endogenous plasma protein levels by generating standard curves in human plasma. Based on their determined plasma protein concentration, they were pooled at approximately 1:1 level to the corresponding endogenous protein in 1µL human plasma. 30 SIS-PrESTs were spiked in an offset between 1 - 20 to the expected endogenous concentrations (**Supplementary Table 1**). The SIS-PrEST pool was dispensed into a 96 well Eppendorf LoBind PCR plate. The plate was vacuum dried at 37 °C for 12 h and stored at -20 °C afterwards [46].

### High load sample digestion

Trypsin from eight different vendors was used to digest four replicates of human plasma and HeLa cell lysate at each 1:50 and 1:20 Enzyme to Protein (E:P) ratio. HeLa cell lysate was only digested in a 1:50 E:P concentration. Each replicate contained 50 µg protein derived from either HeLa cell lysate, 1 µL human plasma or 1 µL human plasma spiked with SIS-PrESTs. Human plasma was 10x diluted in 1x PBS. Vacuum-dried SIS-PrESTs were resuspended in 1x PBS and 10x diluted plasma corresponding to 1 µL plasma was added.

SDC was added to all samples at a final concentration of 0.66 %. Samples were reduced with dithiothreitol (DTT, final concentration 10 mM, 30 min, 56 °C) and alkylated with 2-chloroacetamide (CAA, final concentration 50 mM, 30 min, room temperature (RT) in dark). SDC was diluted to a final concentration of 0.25 % with 1x PBS. The eight trypsins were reconstituted according to vendors recommendations. Only “Sigma 3” was already in solution. All trypsins were diluted to a 0.1 µg/µL concentration in 1x PBS and added in a 1:50 and 1:20 E:P ratio. Samples were incubated at 37 °C shaking over-night before quenching the digestion with trifluoroacetic acid (TFA) to a final concentration of 0.5%.

All samples were desalted through 3-layers of C18 StageTips which were prepared in-house, as previously summarized by Kotol *et al.* [46], [47]. The desalted sample matrix was vacuum-dried at 45 °C and stored at −20 °C prior to MS acquisition.

### Single cell preparation

HEK 293 cells were washed three times with ice-cold PBS to remove any remaining growth media. Cells were resuspended for FACS sorting in fresh, ice-cold PBS at approx. 1e6 cells/ml. Cell sorting was done on a Sony MA900 cell sorter using a 130 µm sorting chip. Cells were sorted at single-cell resolution, into a 384-well Eppendorf LoBind PCR plate containing 1 µl of lysis buffer (80 mM Triethylammonium bicarbonate (TEAB) pH 8.5, 20% 2,2,2-Trifluoroethanol (TFE)). The plate was placed on dry ice for 5 min followed by heating to 95 °C on a PCR machine (Applied Biosystems Veriti 384-well) for 5 min. The cell lysate was digested with Promega 1-3, “Thermo 1” and “Sigma 3” with 2 ng trypsin per cell. For the comparison of trypsin concentrations “Promega 2” was prepared in 0.1 ng, 0.5 ng, 1 ng, 2 ng and 4 ng per cell. 1 uL containing the above stated amount of trypsin was dispensed with an I-DOT One instrument (Dispendix). Samples were incubated at 37 °C overnight and quenched with 1 µL 1 % TFA. Samples were stored at -80 °C prior analysis.

### Liquid Chromatography and Mass Spectrometry setup

SRM assays were run on an Ultimate 3000 nano-LC (Thermo Fisher Scientific) with a TSQ Altis (Thermo Fisher Scientific) MS. An Acclaim PepMap 100 trap column (75 μm × 2 cm, C18, 3 μm, 100 Å, Thermo Scientific) was used together with an analytical PepMap RSLC C18 column (150 μm × 15 cm, 2 μm, 100 Å, Thermo Fisher Scientific) on an EASY-Spray ion source. Samples were loaded onto the trap column at 15 µL/min with 99% solvent A (3% acetonitrile, 0.1% formic acid (FA), H2O) and washed for 0.75 min. Peptides were transferred to the analytical column and separated by a linear gradient of 1-30% solvent B (95% acetonitrile, 0.1% FA) over 29.25 min at a flowrate of 3 µL/min. The linear gradient was followed by three 30 second ramps of 1-95 % Solvent B. During the analysis the column oven was kept at 40 °C, the analytical column was held at 60 °C and the autosampler was kept at 10 °C.

HeLa DDA and Plasma DIA methods were run on a QExactive HF (Thermo Fisher Scientific) with an Ultimate 3000 nano-LC. 2 µg of the sample were injected onto an Acclaim PepMap 100 trap column (75 μm × 150 mm, C18, 3 μm, 100 Å, Thermo Scientific). After flushing the trap column for 3 min at 7 µL/min with 100 % Solvent A the sample is separated on a 40 min linear gradient by an EASY-Spray™ HPLC Columns (75 μm × 250 mm, 2 μm, C18, 100 Å, Thermo Fisher Scientific). The flow was kept at 0.7 µL/min starting at 1 % Solvent B and going to 32 % Solvent B. The linear gradient was followed by three two-minute washes, increasing and decreasing the Solvent B concentration from 1 – 99 %. This was followed by a 9-minute column equilibration. During DIA analysis a full MS scan at 30,000 resolution, AGC = 3e6, 300–1200 m/z, IT = 105 m was performed and followed by 30 10 m/z isolation window scan (30,000 resolution, AGC = 1e6, NCE = 26, IT = 55 ms). For the DDA acquisition a MS1 scan at 60,000 resolution, AGC = 1e6, IT = 205 ms, 300-1200 m/z was performed followed by a top 10 MS2 acquisition with 30,000 resolution, AGC=2e5, IT = 105 ms, 2.0 m/z isolation window and a normalized collision energy (NCE) of 26. Dynamic exclusion was set to 20 sec. During the analysis the column oven was kept at 40 °C, the analytical column was held at 60 °C and the autosampler was kept at 10 °C.

The peptides derived from single cells were separated with the uPAC Neo Low Load analytical column, which was connected to the Ultimate 3000 RSLCnano system according to single-column set-up in the “Ultimate 3000 RSLCnano Standard Application Guide” (page 38) and samples were directly injected from a 384-well plate with a user defined program and a 5uL sample loop. The chromatographic gradient was as follows: Solvent B was increased from 1 to 8.5 % (0 - 4.45min), 8.5 to 12.0 % (4.5 - 5 min), 12 to 25 % (5 - 7 min), 25 to 30 % (7 - 8 min), Solvent B was then increased to 99% and kept constant for 6 minutes (8 – 14.1 min) and dropped to 1% for 4 minutes (14.1 - 18 min). MS spectra were obtained with our recently developed wide window HRMS1 (WISH) – DIA method, that is described in detail here [48], [49]. The FAIMS Pro interface was operated at a compensation voltage of −45 V connected to an Orbitrap Eclipse Tribrid Mass Spectrometer (ThermoFisher Scientific). MS1 scans were performed at 240k resolution with an AGC of 300% and a maximum injection time of 246ms. HCD was used for precursor fragmentation with a NCE of 33 % and MS2 scan AGC target was set to 1000 %. MS2 resolution was set to 240k and an auto injection time with isolation width of 68 m/z. Loop control was set to 2 to insert MS1 for the WISH-DIA modification. This resulted in a MS1 maximum scan cycle time of 1.536 sec.

### SRM Assay Development

The AA sequence of all SIS-PrESTs within this study was *in silico* digested with trypsin using Skyline (**Supplementary Table 1**) [50]. Only peptides between 5-25 AA and no missed cleavages were allowed. Of these peptides all precursors with +2 and +3 charge were included and singly and doubly charged b- and y-ions larger two AA were accepted as transitions. Carbamidomethyl modifications were selected, and R and K were set to ^13^C and ^15^N labeled. SIS-PrESTs were pre-pooled into nine pools that contained between 7 and 25 SIS-PrEST. Pre-pools were based on the most similar endogenous protein concentrations to enable similarly high SIS-PrEST concentrations during assay development. SIS-PrEST pre-pools were digested with trypsin as described above without any plasma background. Each pre-pool digest was analyzed in 3x higher concentration than the expected endogenous level. All *in silico* predicted transitions were searched in this setup with a dwell time over 1 ms and a cycle time of 1 sec for each transition. Precursors that got identified with over 5 transitions were kept and acquired again with a 1 min retention time window (RTW) and a cycle time of 1 sec with a dwell time of over 1 sec. All peptides with a clear peak shape were selected and the top10 transitions accepted while preferring larger product ions and only the peptide precursor with the highest charge state. The collision energy for all peptides was optimized ±5 V from the predicted optimal collision energy. A spectral library was generated for all peptides that passed this assay development. Nine retention time prediction peptides were included to establish an iRT prediction library. 10 µg plasma digest spiked with twice the endogenous level of SIS-PrEST pre-pools was injected with 5 min RTW with a cycle time of 1sec and a dwell time of over 1 ms. All peptides with a clear heavy signal were selected for further assay validation and the five transitions with the highest area under the curve were selected for each precursor. SIS-PrESTs were combined into one final pool containing all SIS-PrESTs from the previously prepared pre-pools. Final SIS-PrEST pool was spiked in a close to 1:1 ratio the endogenous protein into plasma in triplicates. The triplicate digest was injected at a 29.25 min gradient and a 2 min RTW with a cycle time of 1sec and a dwell time of over 1 ms. All peptides that were identified in all replicates with a clear heavy signal were accepted. The CVs of the ratio to standard of each peptide were calculated and the three peptides with the lowest CV of each target protein were selected for the final assay to reduce the number of peptides sufficiently to include all precursors in a single MS method with a gradient of 29.25 min and a cycle time of 1.6 sec with a dwell time of >2 ms and a 2 min RTW. To distribute peptides equally over the full gradient, peptides that were not among the three lowest CVs of a given protein were included if they eluted in in a gradient region with low number of peptides. To assess the quantitative performance of the included peptide LOD, LOQ and linear behavior was established by performing standard curves of SIS-PrESTs in human plasma. The combined SIS-PrEST pool was spiked at 16x the endogenous level into human plasma and serially diluted two-fold in 12 steps. With the previously established SRM method triplicate injection of the standard curve was performed. Only peptides displaying a linear behavior within the standard curves were accepted as quantitative peptides. Other peptides were excluded. LOD and LOQ of each included peptide is reported in the supplementary SRM assay information (See panorama public). The final SRM assay included 122 proteins with 253 peptides in a 35 min MS method with a 29.25 min active gradient and 7 iRT peptides.

### Data Processing

LC-SRM/MS raw data was imported into Skyline and manually assessed. Peptides with 3-5 identified transitions were accepted. The human canonical proteome was set as background proteome to only include proteotypic peptides. A result report was exported from Skyline for further analysis in R. Peptides with a CV of >20 % in >50 % of the different conditions were excluded to reduce quantitative noise in the dataset. Furthermore, peptides with a mean ratio to standard that is 100 times of the 1:1 ratio were excluded to keep high quantitative accuracy. 108 proteins absolute quantified by 223 peptides remained.

LC-DIA/MS raw files from plasma as well as single cells were analyzed using DirectDIA within the Spectronaut 17 environment. Settings were kept at default except, quantification was set to MS1 and trypsin/p was selected as digestion. Minor (Peptide) Grouping quantification was set to be the highest precursors quantity. The human canonical proteome (20401 entries), the pig proteome (UP000008227, 02.04.2023, 46179 entries) and the *Pichia pastoris* proteome (Taxonomy 4922, 02.04.2023, 5257 entries) were provided as FASTA files. All vendors and timepoints were batch randomized for all acquisitions. Due to LC issues occurring in LC-DIA/MS batch two of the plasma samples, this batch was excluded from the analysis based on high TIC variation and lower number of points per peak in MS1. All raw files of batch two are included in the provided rawfiles. All conditions of the single cell data set were searched individually. Output tables were further analyzed in R. Only proteotypic and protein group specific peptides were excepted and precursor charges <2 were removed. For plasma samples MS1- and MS2 quantities below 100 were removed. For single cell samples MS1- and MS2 quantities below 10 and 3 were removed. Single cells with a data completeness below 60 % for the respective condition were excluded from the analysis (12 out of 187 cells). Furthermore, the highest precursor of a given peptides as selected as peptide quantity.

LC-DDA/MS raw files were analyzed using MaxQuant. Trypsin/P was set as enzyme and default settings were kept [51]. The peptide.txt file was imported into R and reversed sequences and potential contaminants were removed.

### Data Availability

DIA, SRM and DDA Raw files together with SRM Skyline documents, spectral libraries and standard curves were uploaded to Panorama Public [52]. The access is currently available upon request but will be public once peer reviewed.

## Results and Discussion

### DDA HeLa Cell digest

HeLa cell lysate was digested in quadruplicate with eight commercially available trypsins at 1:50 (E:P). The digest was acquired with a standard DDA method and searched with Maxquant. No significant differences between the mean protein or peptide levels of the eight trypsins were identified (**Figure 2A**). On average 14.4 % (min. 11.6 %, max. 20.6 %) missed cleavages were detected for all peptides (**Figure 2B**). Here we could identify clear differences between the different trypsin manufacturers. While the Promega trypsins displayed a consistent number of missed cleavages, we could observe substantial variation between the Sigma trypsins. The lowest number of missed cleavages was identified in “Roche 1” (11.6 %) and the highest number of missed cleavages resulted from digestion with “Sigma 2” (20.6 %). With this we concluded that the identified number of peptides seems to be consistent between the eight different trypsins, however, the peptides identified seem to vary between products which could impact the quantification.

**Figure 2.**
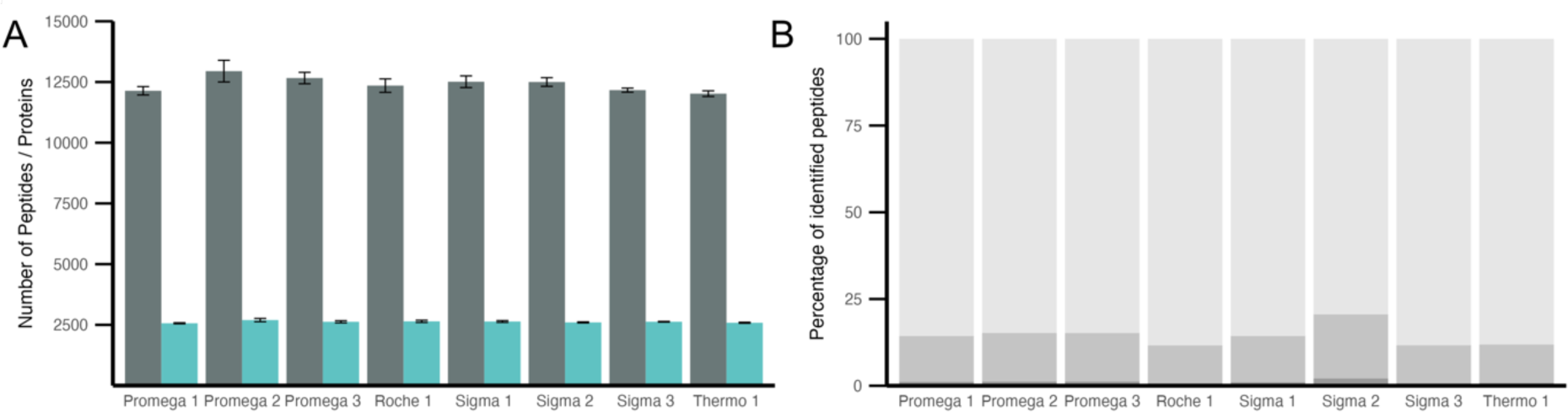
HeLa cell lysate digested with eight commercially available trypsins in a 1:50 E:P. **(A)** Mean identified number of peptides (grey) and proteins (turquoise) out of four digestion replicates. Error bars indicate standard error (SE). No significant differences (<0.05) based on multiple t-test corrected for multiple testing by Bonferroni. **(B)** Percentage of missed-cleavages in peptides identified in ≥3 replicates. 0 (light grey), 1 (medium grey) and 2 (dark grey) missed cleavages for each trypsin shown.

### DIA label free plasma analysis

To further compare the eight commercially available trypsins we assessed their performance in a 1:20 and 1:50 E:P ratio in human plasma in triplicates. Digested plasma samples were acquired using DIA and raw files were searched with Spectronaut. Significant differences between the E:P ratio could be identified for the detected number of peptides but not proteins (**Figure 3A**). Before applying quality control measures as described in the methods section, we quantified 402 ± 15.4 proteins, which were filtered to 317 ± 13.5 to ensure reliable quantitative comparison (**Figure 3B**). Surprisingly, more peptides were identified by all trypsins when using a higher E:P ratio. This difference could also be observed at the protein level for six out of eight trypsins. The different trypsins, delivered for most parts a homogeneous set of identified peptides (**Figure 3C**). The number of unique peptides identified by only one protease was below 30 except for “Sigma 2” which displayed a surprisingly high number of unique peptides. This could potentially be related to the highest number of missed cleavages identified in the DDA-as well as DIA dataset (**Figure 3D**). All trypsins extracted from porcine pancreas displayed a lower number of unique peptides in lower E:P ratio. In contrast, four out of five recombinantly expressed trypsins displayed the opposite behavior. This relationship differed from the missed cleavages which were substantially higher in the 1:50 than the 1:20 E:P. This first DIA comparison of eight trypsins suggested that the peptide and protein numbers between the different proteases show only slight variation. However, an overall trend that suggests higher proteome coverage at a 1:50 E:P which went hand in hand with the number of missed cleaved peptides increasing due to protease concentration (**Supplementary Figure 1**). However, “Sigma 2” displays strong differences to the other proteases, suggesting a bias introduced by this trypsin.

**Figure 3.**
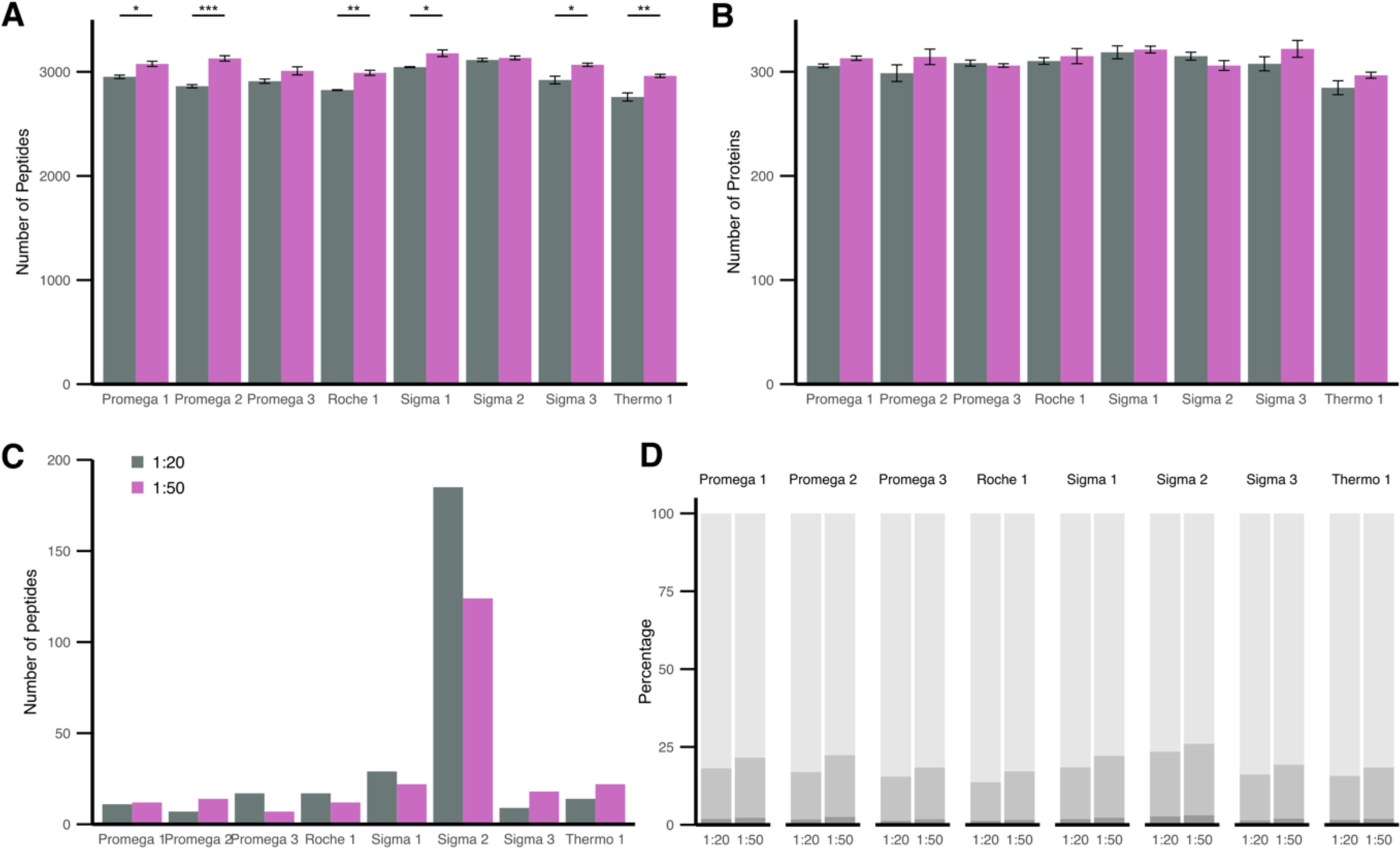
Human Plasma digested with eight commercially available trypsins in a 1:20 (grey) and 1:50 (purple) E:P ratio. **(A)** Mean identified number of peptides out of three digestion replicates. **(B)** Mean identified number of proteins out of three digestion replicates. A-B Error bars indicate SE. Significant differences based on t-test corrected for multiple testing by Bonferroni shown (* p<0.05, ** p <0.01, *** p<0.0001). **(C)** Peptides only identified by respective trypsin (≥50 % of replicates) and not present in other trypsins. **(D)** Percentage of missed-cleavages in peptides identified in ≥50 % of replicates. 0 (light grey), 1 (medium grey) and 2 (dark grey) missed cleavages for each trypsin shown.

To assess the quantitative impact of trypsin we first investigated the proteome depth that could be covered by each protease and the percentage of total identified plasma proteome (361 proteins) covered by all trypsins (**Figure 4A**). While trypsin at 1:20 E:P could identify a mean of 84.1 % ± 3.6 % of the overall identified proteome (361 proteins), 1:50 E:P displayed a trend towards higher proteome coverage in six out of eight trypsins with a mean coverage of 85.7 % ± 2.42 %. “Thermo 1” displayed the overall lowest proteome coverage 76.7 % identified proteome in a 1:20 E:P. This was in line with the previous results that also displayed the lowest number of peptides and proteins for “Thermo 1”.

**Figure 4.**
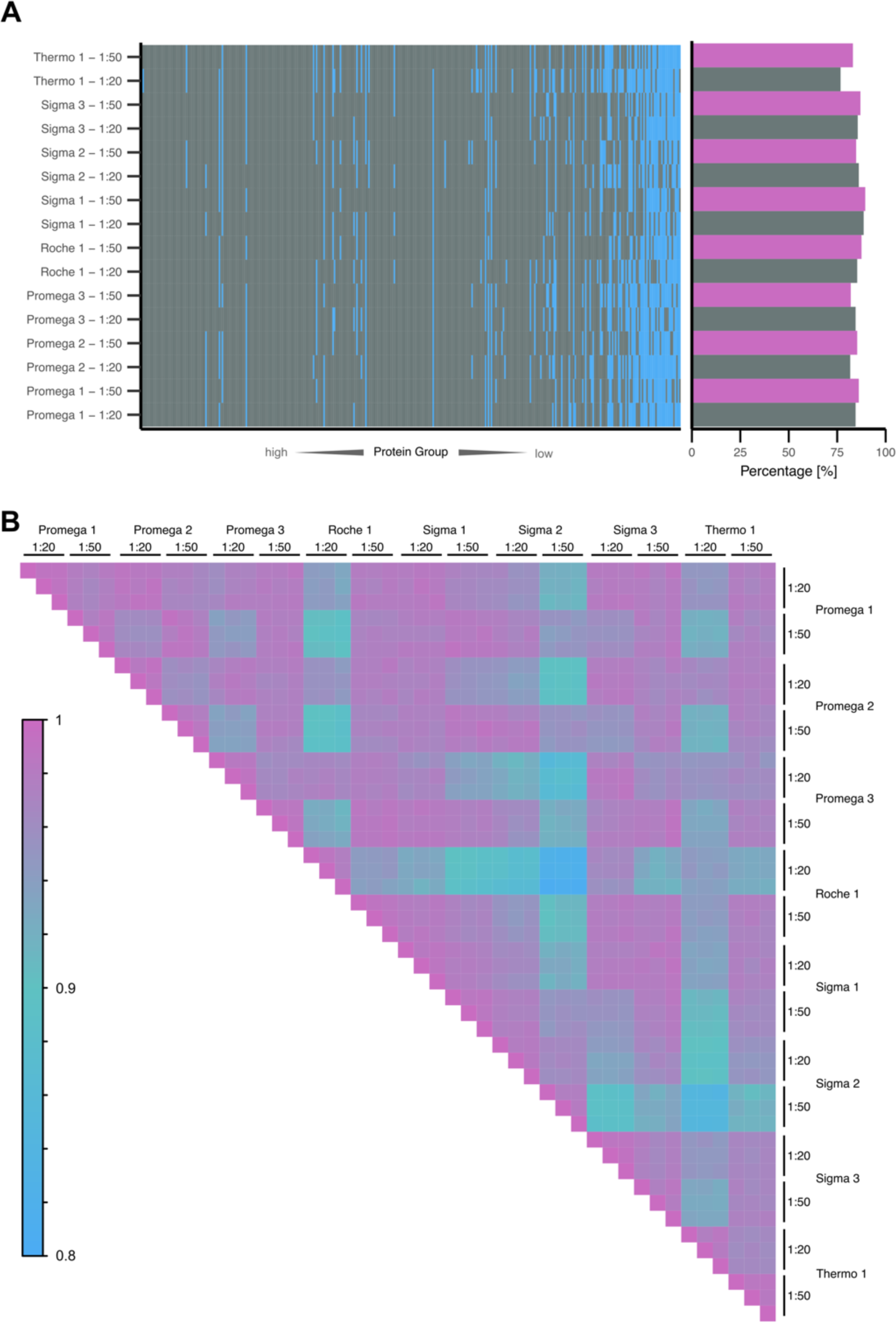
Proteome depth and quantitative correlation of plasma samples acquired in DIA and digested with eight commercially available trypsins in a 1:20 (grey) and 1:50 (purple) E:P ratio. **(A)** All protein Groups identified in the total dataset order by median quantity from highest (left) to lowest (right). Protein groups identified in ≥2 replicates colored grey, protein groups not identified blue. Percentage of identified protein groups of the total number of identified protein groups in the dataset shown to the right. **(B)** Pearson correlation between 1928 quantified peptides in 3 digestion replicates of eight trypsins in two different digestion concentrations. Peptides quantities are log2 scaled.

By Pearson correlation of 1928 peptides quantified in each replicate of the DIA dataset, we assessed the LFQ quantitative performance of each trypsin (**Figure 4B**). Intra-replicate correlations were very high, suggesting that the quantitative impact of sample preparation was neglectable. Besides that, all replicates showed a correlation above 0.8 which was overall in high agreement with the quantitative performance. However, “Sigma 2” displayed lower correlation coefficients at the 1:50 E:P towards other proteases which can be compensated for by a 1:20 E:P ratio. “Roche 1” and “Thermo 1” displayed a lower correlation coefficient at a 1:20 E:P which differed to the 1:50 E:P. Therefore, the protease concentration seemed to impact the quantitative correlation stronger than the choice of protease. However, unique proteases such as “Sigma 2” displayed overall lower agreement with other proteases in terms of quantitative performance.

### SRM SIS Protein based plasma protein quantification

The trypsin concentration seemed to strongly impact the quantitative compatibility of plasma samples analyzed in LFQ DIA data. We further validated these impacts by spiking plasma with heavy labeled protein standards (SIS-PrESTs) and quantified 223 peptides of 108 plasma proteins with absolute precision. The peptide concentration was determined in ratio to standard. We could observe a median CV of 4.4 % between quadruplicate digestion of each trypsin at two concentrations suggesting high technical reproducibility. Overall, a digestion with a 1:20 E:P lead to slightly lower CVs (median 4.08 %) vs 4.8 % median CVs at a 1:50 E:P. Here, we observed the overall lowest CVs for “Sigma 3” (**Figure 5A**). Similar to the DIA dataset, we assessed the quantitative agreement of the different trypsins by Pearson correlation (**Figure 5B**). Overall, very high agreement between the quantified peptides could be observed. Interestingly the impact of trypsin concentration was reduced in comparison to the LFQ DIA quantification. On the other hand, it becomes more apparent that “Sigma 2” displays overall lower quantitative correlation which is supported by all previous findings. The targeted absolute quantification assays did not seem to be as susceptible to the impact of different trypsins and concentrations of trypsin than LFQ DIA sample sets.

**Figure 5.**
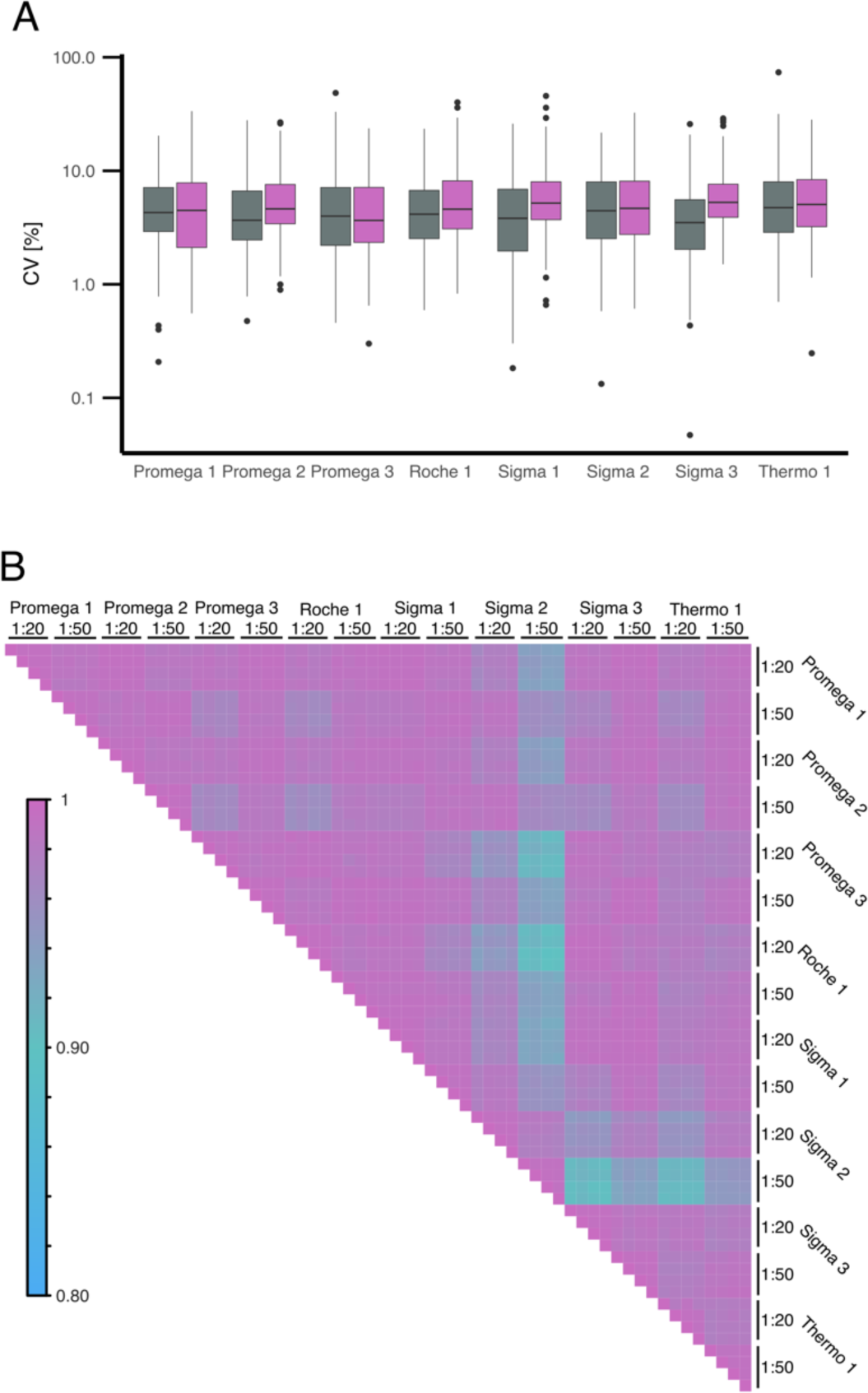
Quantitative variation of absolute quantified peptides based on ratio to standard to SIS-PrEST related to commercially available trypsins and a 1:20 (grey) and 1:50 (purple) E:P ratio. **(A)** CV of 223 peptides quantified in quadruplicates. **(B)** Pearson correlation of log2 ratio to standard between 0.1 and 10 of 223 peptides identified in eight trypsins and two trypsin concentrations.

### Digestion of single cells

To assess whether trypsin is impacting scp-MS in the same manner as high-load samples, 187 single HEK 293 cells were digested with Promega 1-3, “Sigma 3” and “Thermo 1” at 2 ng per cell. While a high degree of technical consistency could be observed for all trypsins, we did observe differences on the quantitative performance of the five different trypsins with “Promega 1”, “Promega 2” and “Sigma 3” showing the highest similarity (**Figure 6A**). The overall variation between sc peptide quantities of different trypsins showed no clear differences, with an average CV between 20.2 % and 28.0 % (**Figure 6B**). Interestingly, the number of peptides identified in single cells varied significantly between two groups of trypsins. “Promega 1” displayed the lowest number of peptides and “Thermo 1” displayed the highest number (**Figure 6C**). This trend did not correspond to the above-described variation observed in high-load samples. As we observed a strong impact of the trypsin concentration on quantitative comparisons, we assessed five trypsin concentrations to evaluate this effect at sc level. To this end, single cells were digested with 0.1 ng, 0.5 ng, 1 ng, 2 ng and 4 ng, using only “Promega 2”. “Promega 2” was chosen based on its widespread use in previous literature. Increasing the protease concentration from 0.1 ng to 0.5 ng boosted the peptide numbers significantly (p = 1.68e-16) (**Figure 6D**). However, for higher concentrations, no clear trend towards further increased peptide numbers could be observed. However, similar to the observations in high load samples, the number of missed cleavages was reduced with increased trypsin concentration (**Figure 6E**). We could observe a mean decrease of missed cleavages of 11.3 % between the highest and lowest concentrations. In summary the choice of trypsin seemed to impact scp-MS experiments differently than high-load samples. However, we could report that scp-MS as well as high load samples displayed lower numbers of missed cleavages with higher trypsin concentrations.

**Figure 6.**
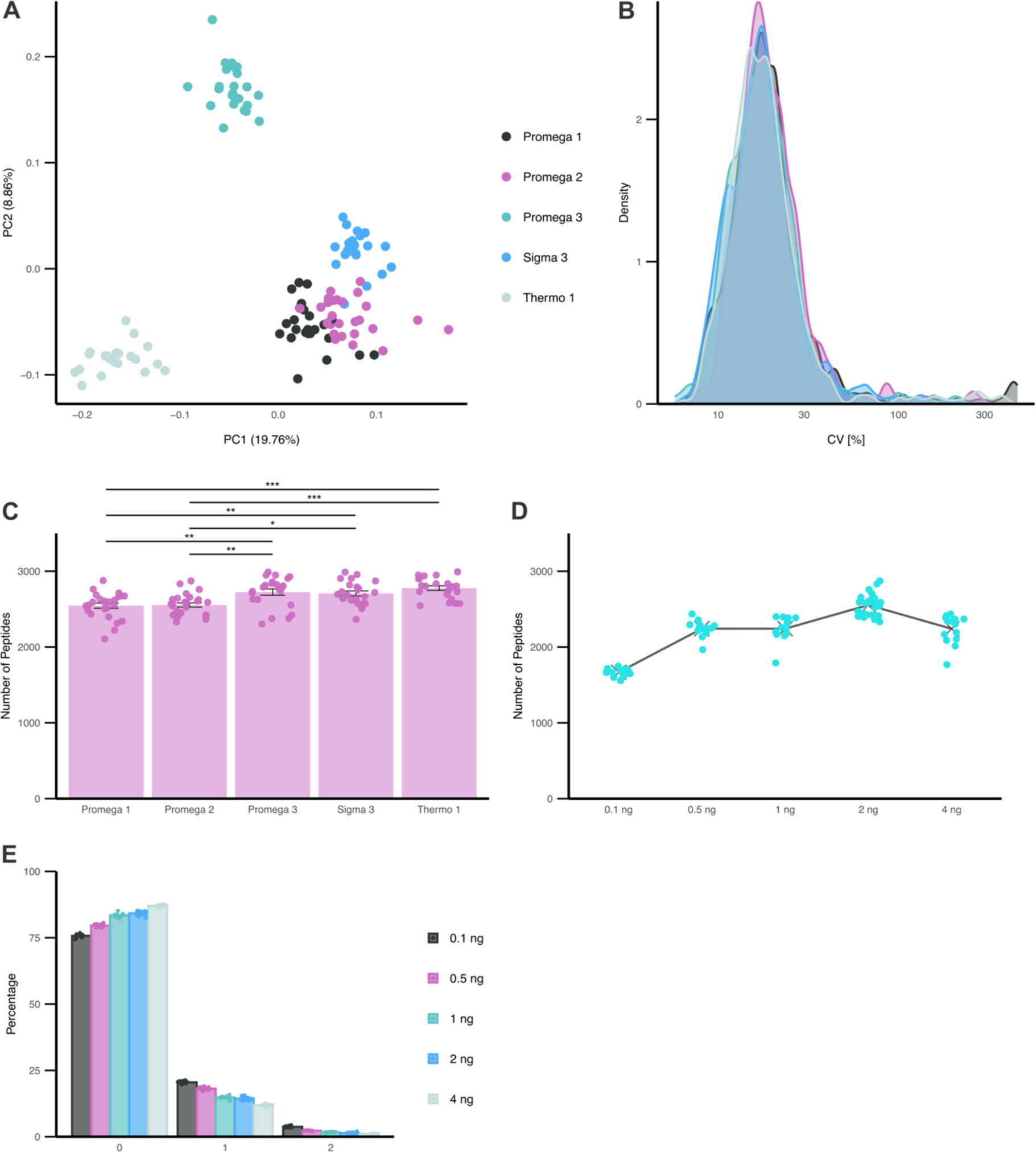
Single HEK293 cell proteomics profiling. **(A)** PCA of single HEK 293 cells digested with five commercially available trypsins. PCA based on peptide level quantities of 610 peptides detected in all cells without imputation. **(B)** CV of 610 peptide quantities identified in all cells. **(C)** Mean identified number of peptides quantified for each commercial trypsin. Each cell highlighted individually. Error bars indicate SE. Significant differences based on t-test corrected for multiple testing by Bonferroni shown (* p<0.05, ** p <0.01, *** p<0.0001). **(D)** Number of peptides quantified in single cells for 0.1 ng, 0.5 ng, 1 ng, 2 ng and 4 ng of “Promega 2”. Sc and total mean shown. **(E)** Percentage of missed cleavages in single cells for 0.1 ng, 0.5 ng, 1 ng, 2 ng and 4 ng of “Promega 2”. Mean of all single cells shown as bar, with replicate cells shown individually.

### The trypsin source could be a significant mediator of trypsin performance

In this work, we compared trypsins which were recombinantly expressed to those extracted from porcine pancreas as described in **Table 1**. To evaluate whether the source of purified trypsin has a significant effect on its performance, we searched the plasma samples acquired in DIA with the respective expression source proteome as background to the human proteome together with the AA sequence of trypsin. Overall, the number of peptides corresponding to the trypsin source made up 14-26 different proteins (**Figure 7A**). In total, it contained 32 porcine- and 35 *Pichia Pastoris* proteins. The number of porcine peptides per trypsin was slightly higher than the number of *Pichia Pastoris* peptides. We furthermore identified between 5 and 15 trypsin-derived specific peptides that were present in both trypsin concentrations of each vendor (**Figure 7B**). These peptides were derived from trypsin itself and not the source of trypsin. The quantity of trypsin-derived peptides was always higher in the 1:20 than the 1:50 assuring correct experimental procedures. However, the intensity of trypsin-derived peptides extracted from porcine pancreas was overall lower than the mean human peptide quantity of the given condition. In comparison, all trypsins expressed in *Pichia Pastoris* except “Sigma 2” display higher trypsin-derived peptide intensities than the mean human peptide quantity. “Sigma 2” shows overall the lowest trypsin-derived peptide quantities in the whole dataset. Trypsin-derived peptide quantities do not necessarily relate to the trypsin amount present during digestion as trypsins could differ in their auto-degradation activity. While “Sigma 2” and “Thermo 1” display the overall lowest quantity of trypsin-derived peptides they had the highest quantitative variations and lowest number of identified peptides and proteins in all high-load sample matrices and acquisition techniques. This could potentially indicate lower overall trypsin concentrations for these two commercially available trypsins. Overall, recombinantly expressed trypsins except “Sigma 2” display a higher quantity of trypsin-derived peptides with lower variation than the porcine counterpart. These findings could provide a possibility to control for trypsin-induced biases in bottom-up proteomics. The trypsin-derived peptide quantities could be used as a measure of digestion efficiency and protease concentration confirmation.

**Figure 7.**
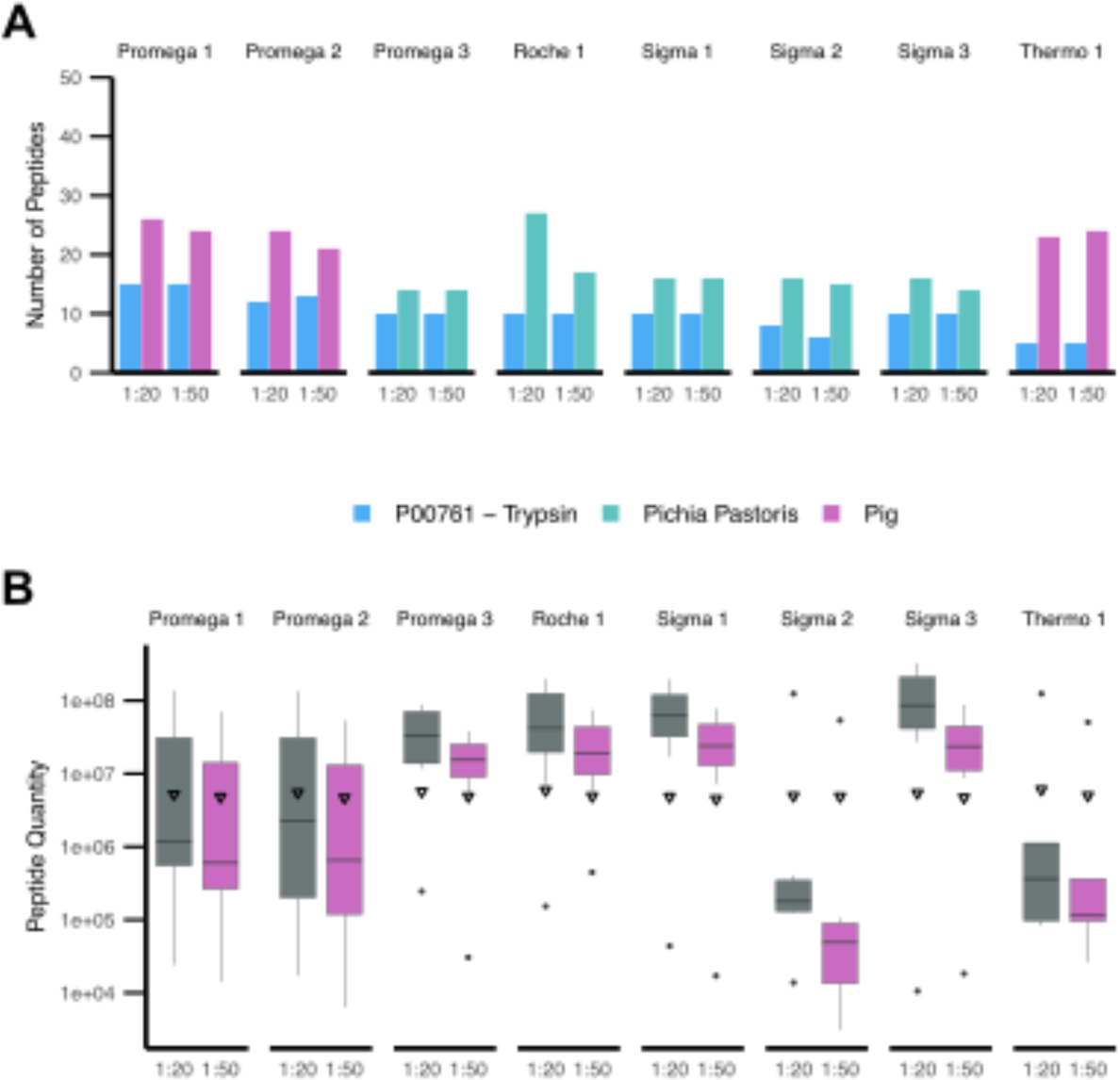
Trypsin related contaminations and autocleavages products identified in human plasma digested with trypsin of eight commercially available vendors and analyzed with DIA. **(A)** Porcine, *Pichia Pastoris* and trypsin specific peptides identified in ≥2 replicates. **(B)** Quantity of all tryptic peptides identified in both 1:20 (grey) and 1:50 (purple) trypsin and in ≥2 replicates. Median peptide intensity of ≥2 replicates of each peptide. Triangle displays mean quantity of all human peptides present in ≥2 replicates.

### Impact of storage time on trypsin performance

Finally, we evaluated the impact of storage time in lyophilized form on “Thermo 1” which was ordered and stored at four time points (0, 7, 12, 14 months) prior to experiments. No significant differences between identified protein or peptide numbers could be identified when analyzing HeLa cell digest acquired in DDA (**Figure 8A**). However, it is worth to note that the number of missed cleavages steadily increased slightly by mean 0.7 % with an increased storage time (**Figure 8B**). This observed increase was however not as prominent as the observed variation between different trypsins, thereby suggesting merely a minor impact which should be further validated. The quantitative performance in LFQ DIA data showed no significant variation from mean (**Figure 8C**). By further looking into single peptide variation over time, we explored the ratio-to-standard variation of peptides in the SRM assay (**Figure 8D**). 68.6 % of the peptides displayed a CV below 5 % over the digestion with “Thermo 1” stored over different periods of time (**Figure 8E**). Only 4.9 % of the peptides displayed CVs over 10 % in their ratio to standard. This indicates that the storage time of “Thermo 1” did not seem to impact the quantitative performance of the trypsin.

**Figure 8.**
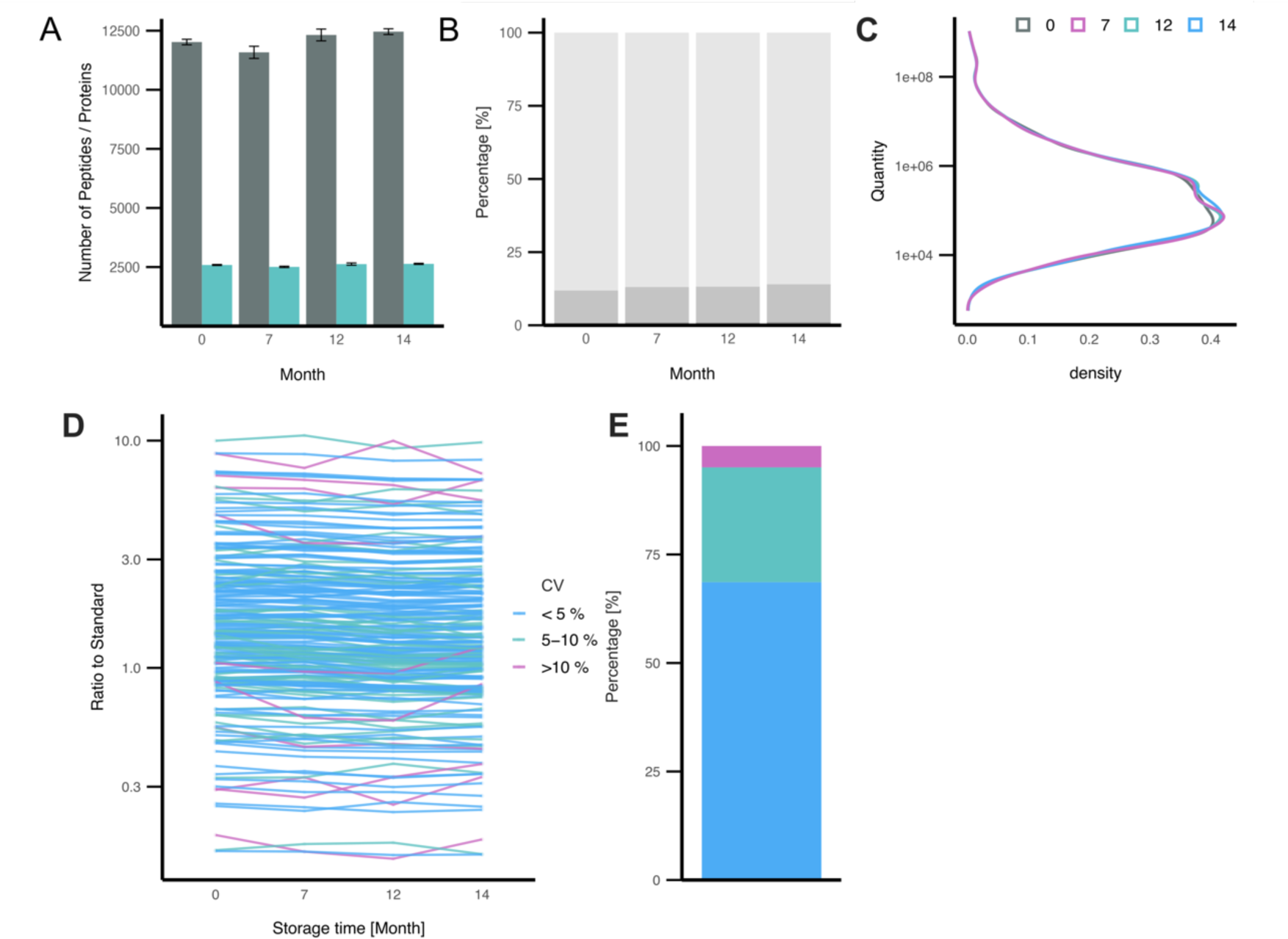
Lyophilized **“**Thermo 1” trypsin was ordered at four timepoints over the course of 14 month. **(A)** HeLa cell digest acquired in DDA. Mean identified number of peptides (grey) and proteins (turquoise) out of four digestion replicates. Error bars indicate SE. No significant differences (>0.05) based on multiple t-test corrected for multiple testing by Bonferroni. **(B)** HeLa cell digest acquired in DDA. Percentage of missed-cleavages in peptides identified in ≥3 replicates. 0 (light grey), 1 (medium grey) and 2 (dark grey) missed cleavages for each trypsin shown. **(C)** Density plot of peptide quantities identified 1:50 digestion in ≥2 replicates in DIA acquisition. Each density plot corresponds to one storage time. **(D)** Ratio to standard of absolute quantified peptides in SRM for each trypsin storage timepoint. Each line corresponds to one peptide. For visualization purposes only peptides with a ratio to standard between 0.1 and 10 included. Color corresponds to the CV of the ratio to standard over the four timepoints (blue <5 %, turquoise 5-10 %, violet >10 %). **(E)** Percentage of peptides shown in D that display CV as described.

## Conclusions

In this study, we extensively investigated the impact of eight commercial trypsins on commonly used bottom-up proteomics workflows, across a range of samples. Besides the trypsin itself, different protease concentrations, sample matrices, instrument acquisition techniques and protease storage times were investigated. The resulting dataset allowed us to unravel some of the technical impacts proteases have in bottom-up proteomics, while the field is transitioning further into large-scale clinical- and biomedical research applications. In high-load samples, we observed low variation between the trypsins regarding the pure number of proteins or peptides identified in each sample and in terms of proteome depth. Differences only became apparent when looking into the unique peptides that each protease generates and the number of missed cleavages. From a quantitative standpoint, the majority of the trypsins performed similarly in each given concentration except for “Sigma 2”. In scp-MS, however, the choice of trypsin seemed to display stronger impacts on the number of peptides and their quantitative performance. On the other hand, trypsins displayed similar technical variation between cells, and minor cell heterogeneity was apparent irrespective of which trypsin was used. Unexpectedly, the concentration of trypsin had a much more drastic impact on the quantitative and qualitative performance of trypsin both in plasma, bulk and scp-MS. A higher trypsin concentration tended to result in more comparable protein quantitation, besides also giving rise to lower numbers of missed cleavages. Surprisingly however, the higher trypsin concentration did not result in a higher number of identified proteins or peptides for plasma and HeLa bulk digest. In scp-MS we could observe increased numbers of peptides which plateaued towards higher concentrations.

Our results indicated that trypsins with higher quantitative variation (e.g. “Sigma 2”, “Thermo 1”) also showed the lowest quantity of trypsin-derived peptides in the samples. This suggests that the product specific variability might not derive from the enzyme itself but rather its concentration or activity. As all trypsins display a rather low number of contaminating peptides identified in plasma samples, this should not be a specific criterion for trypsin selection. However, all recombinantly expressed trypsins except “Sigma 2” displayed higher concordance between their trypsin peptide quantities. This could suggest that recombinantly expressed trypsins might be easier to interchange with one another than trypsins from different sources. However, further evidence is needed. Furthermore, we see indications that the monitoring of trypsin peptides in datasets could provide an opportunity to compensate for trypsin related biases in bottom-up proteomics.

When investigating the effects of long-term storage of a single trypsin, we could not observe any strong storage time related impacts on quantitative levels. This suggests that long term trypsin storage might be a way to compensate for trypsin related quantitative variations in bottom-up proteomics analysis of large cohorts. With that higher reproducibility might be achieved by storing the required amount of trypsin in one batch rather than buying new batches on an on-going basis.

When considering the large number of samples that are processed in clinical tests or current cohorts the price of trypsin per sample is a substantial consideration. Our data suggests that the protease itself does not show substantial quantitative differences between vendors for most parts, but the protease concentration should be further investigated. While trypsin concentration is a factor to consider when preparing larger cohorts, it seems advisable to choose a commercial trypsin with reproducible trypsin peptide quantities. This would potentially reduce the concentration bias. As storage time does not seem to impact the trypsin performance to a strong degree, it might be useful to reduce trypsin variability by preordering large batches of trypsin produced in one batch to compensate for trypsin concentration variation. However, to validate the relevance of these results for specific experimental setups, these variations should be assessed for each specific trypsin and digestion condition. It must be noted that the digestion conditions have previously been described as a major impact on the protease digestion performance and the presented findings here might not necessarily translate to other digestion protocols [20].

In summary we have provided a large dataset comparing eight commercially available trypsins on three different biological samples that are frequently used in biomedical research and will find future applications in clinical tests. We observed minor differences in performance between trypsins, with the trypsin concentration having the largest effect on the quantitative performance of bottom-up experiments. Notably, the use of recombinant protein standard for absolute quantification provided a way to reduce the protease related bias in comparison to LFQ. This could be a key aspect to consider while the field moves further into clinical applications. Therefore, we suggest paying special attention to the choice of trypsin when studying large cohorts with bottom-up proteomics and validate its reproducibility. Certain trypsins might be interchangeable with one another which is an important aspect to consider when moving forwards into clinical applications. However, further studies in this regard are needed.

## Acknowledgments

We thank Christian Gnann and Sarah Line Skovbakke for providing cells for the conducted experiments. Furthermore, we thank the protein factory of the Human Protein Atlas program and the Science for Life Laboratory for their contributions. Ulrich auf dem Keller acknowledges the Novo Nordisk Foundation Young Investigator Award (NNF16OC0020670 to U.a.d.K).

