## SupplementaryFigure1 for "Assessing the role of trypsin in quantitative plasma- and single-cell proteomics towards clinical application"

* Corresponding author

[+] Contributed equally to this work

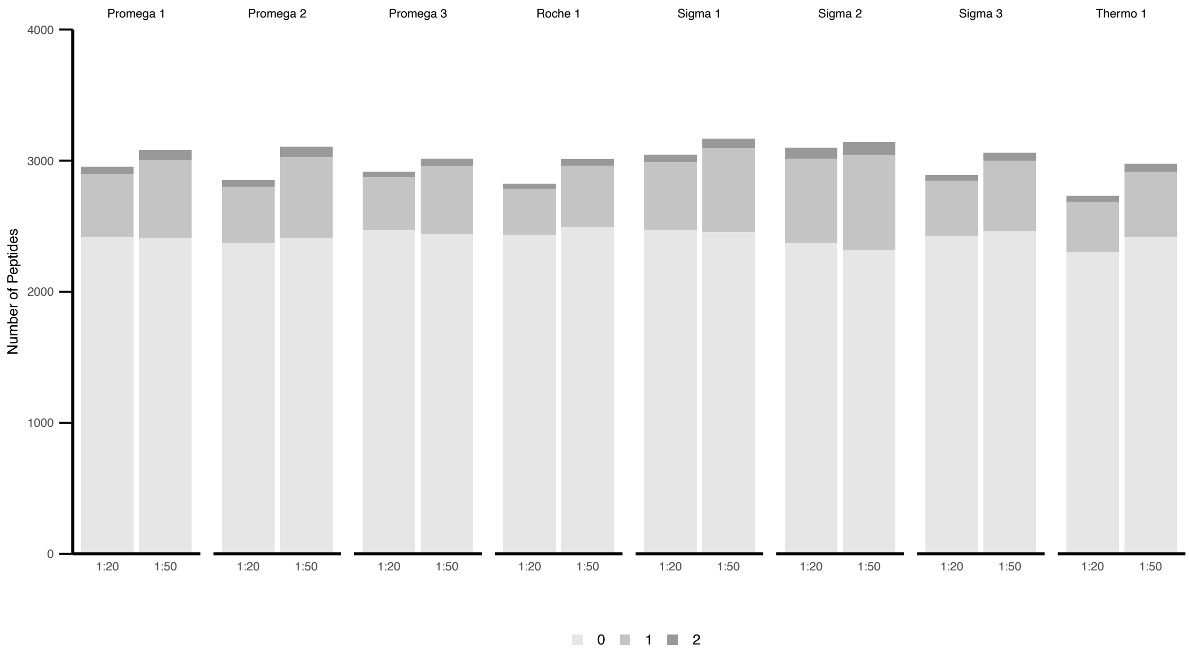


**Supplementary Figure 1.** Human Plasma digested with eight commercially available trypsins in a 1:20 and 1:50 E:P ratio. **(A)** Median number of identified peptides and respective missed cleavages out of three digestion replicates. Missed cleavages are displayed in grayscale.
